# *Trypanosoma brucei* J protein 2 functionally cooperates with the cytosolic Hsp70.4 and Hsp70 proteins

**DOI:** 10.1101/641704

**Authors:** Stephen J. Bentley, Aileen Boshoff

## Abstract

The etiological agent of African trypanosomiasis, *Trypanosoma brucei*, has been identified to possess an expanded and diverse group of heat shock proteins, that have been implicated in cytoprotection, differentiation, and subsequently progression and transmission of the disease. Heat shock protein 70 is a highly conserved and ubiquitous molecular chaperone that is important in maintaining protein homeostasis in the cell. Its function is regulated by a wide range of co-chaperones; and inhibition of these functions and interactions with co-chaperones are emerging as potential therapeutic targets for numerous diseases. This study sought to biochemically characterize the cytosolic Hsp70 and Hsp70.4 proteins and to investigate if they form a functional partnership with the Type I J-protein, Tbj2. The cytosolic localisation of the proteins was confirmed by accessing the TrypTag endogenous tagging microscopy database. Expression of TbHsp70 was shown to be heat inducible, whilst TbHsp70.4 was constitutively expressed. The basal ATPase activities of TbHsp70.4 and TbHsp70 were stimulated by Tbj2. It was further determined that Tbj2 forms a functional partnership with TbHsp70 and TbHsp70.4 as the J-protein was shown to stimulate the ability of both proteins to mediate the refolding of chemically denatured β-galactosidase. This study provides further insight into this important class of proteins which may contribute to the development of new therapeutic strategies to combat African Trypanosomiasis.

## Introduction

African trypanosomiasis, a neglected tropical disease, afflicts humans as well as domestic and wild animals, and has a detrimental impact on socioeconomic development in sub-Saharan Africa [1]. There is a need for the development of more effective and safer chemotherapies to treat the disease, due to existing drug toxicity, growing parasite resistance and the lack of a vaccine [2]. The Hsp70/J-protein chaperone machinery has been implicated to play an integral role in the development, differentiation, and survival of protozoan parasites, as they transition through the various stages of their life cycle [3]. In *Leishmania* and *Trypanosoma cruzi*, heat shock proteins have been shown to play an essential role in stress-induced stage differentiation and are important for disease progression and transmission [4–5], making this protein family an attractive chemotherapeutic target. The completion of the *Trypanosoma brucei* (*T. brucei*) genome has expedited transcriptome and proteome analyses and revealed that the extracellular parasite has an expanded and diverse Hsp70 and J-protein complement, with the parasite possessing cytosolic Hsp70 members that display atypical Hsp70 features [6].

The Hsp70 protein family is ubiquitous and plays an integral role in protein quality control and maintaining protein homeostasis under normal and stressful conditions [7–8]. Evolution has given rise to multiple homologous *Hsp70* genes, with Hsp70 members found in all the major subcellular compartments within the cell [9–10]. The cytosol of eukaryotic cells has been shown to possess two major Hsp70 isoforms, a stress-inducible (Hsp70) and constitutively expressed (Hsc70) form [9]. The structure of eukaryotic cytosolic Hsp70s are highly conserved and are typically comprised of an N-terminal nucleotide binding domain and a C-terminal domain with a substrate binding domain (SBD) and a 10 kDa α-helical domain with a conserved EEVD motif [11–12].

Hsp70 functions as either an ATP-dependent refoldase, which entails folding nascent and unfolded polypeptides into their respective native states; or displays ATP-independent holdase activity, which involves binding to unfolded polypeptide aggregates and retrieving them back into solution [13]. The diversity of the roles performed by Hsp70s are driven by a cohort of proteins known as co-chaperones [7], including J proteins, nucleotide exchange factors (NEFs), and tetratricopeptide repeat (TPR) domain containing proteins [14]. One of the most important classes of Hsp70 co-chaperones are J proteins. Type I J-proteins are comprised of an N-terminal J-domain with a conserved HPD motif, a glycine-phenylalanine rich region, four zinc finger motifs (zinc finger domain) and a C-terminal peptide binding domain [15–18]. Type II J-proteins have a glycine-methionine rich region instead of the zinc finger domain [19]. Both type I and type II J-proteins serve to bind substrate polypeptides and target them to Hsp70 for refolding [19]. Type III J-proteins only possess the J-domain, however, it is not necessarily located at the N-terminal [19]. Type IV J-proteins also possess the J-domain, but the HPD motif is non-conserved or absent [20].

A recent *in silico* investigation revealed that the *T. brucei* genome was found to encode 12 members of the Hsp70 superfamily, with 8 members from the Hsp70/HSPA family and 4 Hsp110/HSPH family members [6]. The same study identified 67 putative J-proteins, with 5 type I J-proteins [6]. Phenotypic knockdown of *T. brucei* genes using RNAi, conducted by Alsford and colleagues [21], demonstrated that the Hsp70/J-protein machinery plays a prominent role in trypanosome biology, as the loss of certain members of these protein families impacted the survival and fitness of the parasite at various stages of its life cycle. Of significance to this study are the cytosolic TbHsp70 and TbHsp70.4 and their interactions with the Type I J-protein Tbj2.

It has been proposed that TbHsp70 is cytosolic and a vital component of the heat shock response as it plays a key role by providing cytoprotection against cellular stress [6]. The Hsp70 orthologue in *Leishmania chagasi* was been linked to the parasite’s resistance to macrophage-induced oxidative stress [22], and has been shown in several *Leishmania* spp. to be linked to parasite’s resistance to pentavalent antimonial treatment, as it induces Hsp70 expression which provides stress tolerance against the drug [23–24]. Based on phylogeny, TbHsp70.4 was found to form a distinct Hsp70/HSPA group unique to kinetoplastid parasites, with no obvious mammalian orthologues and a divergent C-terminal EEVD motif [6]. The Hsp70.4 orthologue in *Leishmania major* was shown to be cytoplasmic and constitutively expressed [25]. Tbj2 was identified to be an essential stress inducible Type I J-protein, as knockdown via RNAi revealed a severe growth defect and it was shown to reside in the parasite cytosol, [26]. Tbj2 was determined to possess holdase activity when chemically denatured rhodanese and thermally aggregated MDH were used as substrates [27].

Few TbHsp70/J-protein interactions have been biochemically characterised. This study aimed to investigate a potential functional partnership between TbHsp70 and TbHsp70.4 with the co-chaperone Tbj2. The expression of TbHsp70 was determined to be heat inducible, whereas TbHsp70.4 was constitutive. Tbj2 stimulated the ATPase activities of both TbHsp70 and TbHsp70.4. TbHsp70 and TbHsp70.4 were both demonstrated to suppress the aggregation of thermally induced MDH in a dose dependent manner, and this was further enhanced by Tbj2. Furthermore, Tbj2 stimulated the refolding abilities of the Hsp70s of chemically denatured β-galactosidase. Tbj2 does form a functional partnership with both TbHsp70 and TbHsp70.4 and is part of the functional chaperone network that has been implicated in the proliferation and growth of parasitic cells. An increase in knowledge will enhance our comprehension of this important class of proteins and improve our understanding of the biology of the parasite, which may contribute to the development of therapeutic strategies to combat African Trypanosomiasis.

## Materials and Methods

### The pQE2-TbHsp70 expression vector

The codon optimized coding sequence for expression of TbHsp70 (TriTrypDB accession number: Tb927.11.11330) in *Escherichia coli* was synthesized and supplied by the GenScript Corporation (Piscataway, New Jersey, U.S.A.). The TbHsp70 coding region was inserted into the pQE2 expression vector (Qiagen, U.S.A.) using *Nde*I and *Hind*III restriction sites. The integrity of the pQE2-TbHsp70 vector which was used to express the N-terminal His-tagged TbHsp70 was verified by restriction analysis and DNA sequencing.

### Heterologous expression and purification of TbHsp70 and TbHsp70.4

*E. coli* XL1 Blue cells transformed with either pQE2-TbHsp70 or pQE2-TbHsp70.4 were grown at 37°C in 2x YT medium supplemented with 100 μg/ml ampicillin and grown to mid-logarithmic phase (A_600_ 0.4–0.6). Protein production was induced by the addition of 1 mM IPTG (isopropyl-β-D-thiogalactopyranoside), and the bacterial cultures were incubated at 37°C for 3 hours for TbHsp70 and 1 hour for TbHsp70.4. Bacterial cells expressing TbHsp70 or TbHsp70.4 were harvested by centrifugation (10 000 g; 15 min; 4 °C) and the cell pellet was resuspended in lysis buffer (100 mM Tris-HCl, pH 7.5, 300 mM NaCl, 20 mM imidazole, 1 mM PMSF, 1 mg/ml lysozyme), allowed to stand for 30 min at room temperature and then frozen at −80°C overnight. The cells were then thawed on ice and sonicated at 4°C. The resulting lysate was cleared by centrifugation (13 000 g, 40 min, 4°C) and the supernatant was incubated with cOmplete His-tag purification resin (Roche, Germany) and allowed to bind overnight at 4°C with gentle agitation. The resin was then pelleted by centrifugation (4500 g; 4 min) to remove unbound proteins and washed three times using native wash buffer (100 mM Tris-HCl, pH 7.5, 300 mM NaCl, 50 mM imidazole, 1 mM PMSF) to remove non-specific contaminants. The bound protein was eluted three times by re-suspending the resin in elution buffer (10 mM Tris-HCl, pH 7.5, 300 mM NaCl, 750 mM imidazole). The eluted proteins were extensively dialysed using SnakeSkin dialysis tubing (Pierce-MWCO 10,000; Thermo Scientific, USA) into either dialysis buffer (DB; 10 mM Tris, pH 7.5, 100 mM NaCl, 0.5 mM DTT, 10% (v/v) glycerol, 50 mM KCl, 2 mM MgCl_2_) or into the appropriate assay buffer for functional studies and then subsequently concentrated against PEG 20000 (Merck, Germany). The protein yield was estimated using the Bradford assay (Sigma-Aldrich, USA) with BSA as the standard. SDS-PAGE (10%) and western analysis using mouse monoclonal anti-His primary antibody and HRP-conjugated goat anti-mouse IgG secondary antibody (Santa Cruz Biotechnology, USA) were conducted to assess the expression and purification of the recombinant proteins. TbHsp70 and TbHsp70.4 protein expression in *E. coli* was also confirmed by western blot using rabbit-polyclonal anti-TbHsp70 and rabbit-polyclonal anti-TbHsp70.4 respectively. HRP-conjugated goat anti-rabbit (Santa Cruz Biotechnology Inc., U.S.A.) was used as the secondary antibody. Imaging of the protein bands on the blot was conducted using the ECL kit (Thermo Scientific, USA) as per manufacturer’s instructions. Images we captured using the ChemiDoc Imaging system (Bio-Rad, USA).

### Purification of Tbj2

Recombinant N-terminal His-tagged Tbj2 was purified under native conditions from *E. coli* BL21 (DE3) cells as previously described [27]. Samples were dialysed in DB or into the appropriate assay buffer for functional studies.

### Investigation of heat-induced expression of TbHsp70 and TbHsp70.4 in *T. b. brucei* Lister 927 V221 parasites

Wild type *T. b. brucei* Lister 927 v221 bloodstream form lysates (10^6^ cells/ml) were used for the heat stress inducibility experiment. Bloodstream form *T. b. brucei* Lister 927 v221 strain trypanosome parasites were cultured in filter sterilized complete Iscoves Modified Dulbeccos Media (IMDM) based HM1-9 medium (IMDM base powder, 3.6 mM sodium bicarbonate, 1 mM hypoxanthine, 1 mM sodium pyruvate, 0.16 mM thymidine, 0.05 mM bathocuprone sulphate acid, 10% (v/v) heat inactivated Foetal Bovine Serum, 1.5 mM L-cysteine, 0.2 mM β-mercaptoethanol, pH 7.5) in a humidified chamber at 37°C with an atmosphere of 5% CO_2_. Separate 25 ml culture of cells were exposed to heat shock at 42°C for a period of 120 min in plugged flasks, allowing no entry of CO_2_. A control experiment was performed under the same conditions maintaining the temperature at 37 °C. Cell lysates were harvested by centrifugation at 800 g for 10 min, washed twice in 1x PBS buffer, and repelleted prior to resuspension in SDS-PAGE loading buffer to a final cell count of 5 × 10^5^ cells/μl. The lysates (5 × 10^6^ cells per lane) were resolved on a 10% SDS-PAGE gel. Differences in TbHsp70 and TbHsp70.4 protein expression were detected using rabbit-polyclonal anti-TbHsp70 and rabbit-polyclonal anti-TbHsp70.4 respectively, and goat anti-rabbit IgG HRP-conjugated secondary antibody (Santa Cruz Biotechnology Inc., U.S.A.) in subsequent western analysis. Actin was also probed as a loading control using mouse monoclonal anti-actin antibody and HRP-conjugated goat anti-mouse IgG secondary antibody (Santa Cruz Biotechnology Inc., U.S.A.). Images were acquired using the ChemiDoc Imaging system (Bio-Rad, USA), and densitometric analyses of bands were conducted using the Image Lab v5.1 built 8 (Bio-Rad, USA).

### Sub-cellular localisation of TbHsp70 and TbHsp70.4

Images for the sub-cellular localisation of TbHsp70 (Tb927. 11.11330) and TbHsp70.4 (Tb927.7.710) were kindly sourced from Richard Wheeler from the TrypTag high-throughput microscopy database [28].

### MDH aggregation suppression assay

The capacity of TbHsp70 and TbHsp70.4 to suppress the thermally-induced aggregation of the model substrate, malate dehydrogenase (MDH) from porcine heart (Sigma-Aldrich, U.S.A.), alone or in the presence of Tbj2 was adapted from [29]. Briefly, the reaction was initiated by adding 0.72 μM MDH to varying concentrations of the *T. brucei* Hsp70s in assay buffer (50 mM Tris-HCl pH 7.4, 100 mM NaCl) either alone or in combination with Tbj2. The reaction was then heated at 48 °C for an hour. After incubation, the samples were centrifuged at 13 000 g for 10 minutes to separate the soluble and insoluble fractions. These fractions were analysed using 10% SDS-PAGE, and subsequently quantified using densitometric analysis using the Image Lab v5.1 built 8 (Bio-Rad, USA). A negative control of BSA (0.75 μM) was added to MDH in assay buffer was conducted to illustrate that MDH aggregation suppression was due to the holdase function of the *T. brucei* molecular chaperones and not simply due to the presence of a second protein in the assay system. As a control, the aggregation of the chaperones was monitored in the assay buffer without MDH. Each assay was conducted in triplicate and three independently purified batches of proteins were used.

### ATPase activity assay

The determination of the basal ATPase activity of TbHsp70 and TbHsp70.4 were performed using a high throughput colorimetric ATPase assay kit (Innova Biosciences, U.K.). Summarily, the method allows the quantification of the inorganic phosphate (Pi) released from ATP hydrolysis by an enzyme. Briefly, the Hsp70s were prepared in ATPase assay buffer (100 mM Tris-HCl, 7.5, 2 mM MgCl_2_) and incubated with varying ATP concentrations (0-2 mM) for 1 hour at 37 °C. The samples containing Pi hydrolysed from ATP were incubated with the PiColorLock™ solution, which is a malachite green dye solution that in the presence of Pi changes absorbance due to the generation of molybdate-phosphate complexes. The absorbance was measured at 595 nm using a Powerwave 96-well plate reader (BioTek Instruments Inc., U.S.A.), and absorbance values were converted to phosphate concentrations using a standard curve of absorbance vs. phosphate concentration based on a set of Pi standards provided by the supplier assayed along with the samples. All samples were corrected for spontaneous breakdown of ATP observed in a control experiment in the absence of protein. A Michaelis-Menten kinetic plot was constructed and used to determine the kinetic specific activity (*V*max) and *K*m for the basal ATPase activity of TbHsp70 and TbHsp70.4. A non-linear regression curve was fitted to the Michaelis-Menten plot using GraphPad Prism® (v. 7.0; San Diego, CA, U.S.A.) software. The assay was conducted in triplicate on three independently purified batches of proteins.

To investigate the modulatory effect of Tbj2 on the basal ATPase activity of the *T. brucei* Hsp70s, the molecular chaperones at indicated concentrations were prepared in ATPase assay buffer (100 mM Tris-HCl, 7.5, 2 mM MgCl2) and incubated with 1 mM ATP for 1 hour at 37 °C. The colour development and absorbance measurement procedures were conducted as previously described. All samples were corrected for spontaneous breakdown of ATP observed in a control experiment in the absence of protein. Additional control of the respective boiled Hsp70 protein was used to cater for the spontaneous hydrolysis of ATP. Any background ATP hydrolysis observed for Tbj2 was corrected for by subtracting this activity from the reactions containing these proteins. The modulatory effect on the ATPase activity of the *T. brucei* Hsp70s was represented as fold change with the basal ATPase activity of the Hsp70s taken as 1. The assay was conducted in triplicate on three independently purified batches of protein.

### The β-galactosidase refolding assay

Investigation of the ability of the *T. brucei* Hsp70s to refold chemically denatured β-galactosidase alone or in combination with Tbj2 was carried out as previously described [30]. The catalytic hydrolysis activity of the refolded β-galactosidase using 2-Nitrophenyl β-D-galactopyranoside (ONPG) as a chromogenic substrate [31] is measured in relation to native β-galactosidase and used as means to determine the refoldase activity of Hsp70. After denaturation of β-galactosidase (Sigma-Aldrich, U.S.A.) for 30 minutes at 30 °C in denaturation buffer (25 mM HEPES, pH 7.5, 5 mM MgCl_2_, 50 mM KCl, 5 mM β-mercaptoethanol, and 6 M guanidine-HCl), denatured β-galactosidase to a final concentration of 3.4 nM was diluted 1:125-fold into refolding buffer (25 mM HEPES pH 7.5, 5 mM MgCl_2_, 50 mM KCl, 2 mM ATP, and 10 mM DTT) supplemented with the *T. brucei* Hsp70s alone or in the presence of Tbj2 at indicated concentrations, and incubated for 2 hours at 37 °C. Reactions containing 1.6 μM BSA (no molecular chaperones) with native and denatured β-galactosidase was conducted to serve as positive and negative controls respectively. The activity of β-galactosidase was measured at various time points by mixing 10 μl of each refolding reaction with 10 μl of ONPG, followed by incubation at 37°C for 15 min. Assays were terminated by the addition of 0.5 M sodium carbonate and the absorbance of each sample was measured spectrophotometrically at 412 nm. The percentage refolding activity is calculated relative to the activity of native β-galactosidase. The assay was conducted in triplicate on three independently purified batches of protein.

## Results and Discussion

### Recombinant protein production and purification

TbHsp70 and TbHsp70.4 recombinant proteins were both expressed in *E. coli* XL1 Blue cells and the His-tagged proteins were successfully purified using nickel affinity chromatography (Fig 1). Both *T. brucei* Hsp70 proteins were eluted as species of approximately 71 kDa (Fig 1A-B). However, for TbHsp70.4 protein species of lower molecular weights were also produced in *E. coli* along with the full-length recombinant protein (Fig 1B). Anti-TbHsp70.4 and anti-His antibodies were both able to recognise the full-length TbHsp70.4 recombinant protein and the lower molecular weight species as confirmed by western analysis (Fig 1B). These protein species were either N-terminally His-tagged truncated versions of the full-length TbHsp70.4 protein or products of incomplete synthesis. Despite this, TbHsp70.4 was purified by nickel affinity chromatography as full-length protein (Fig 1B). Recombinant Tbj2 was expressed in *E. coli* BL21 (DE3) cells, and subsequently purified using nickel affinity chromatography as previously described [27] (Fig S1A-B).

**Fig 1.**
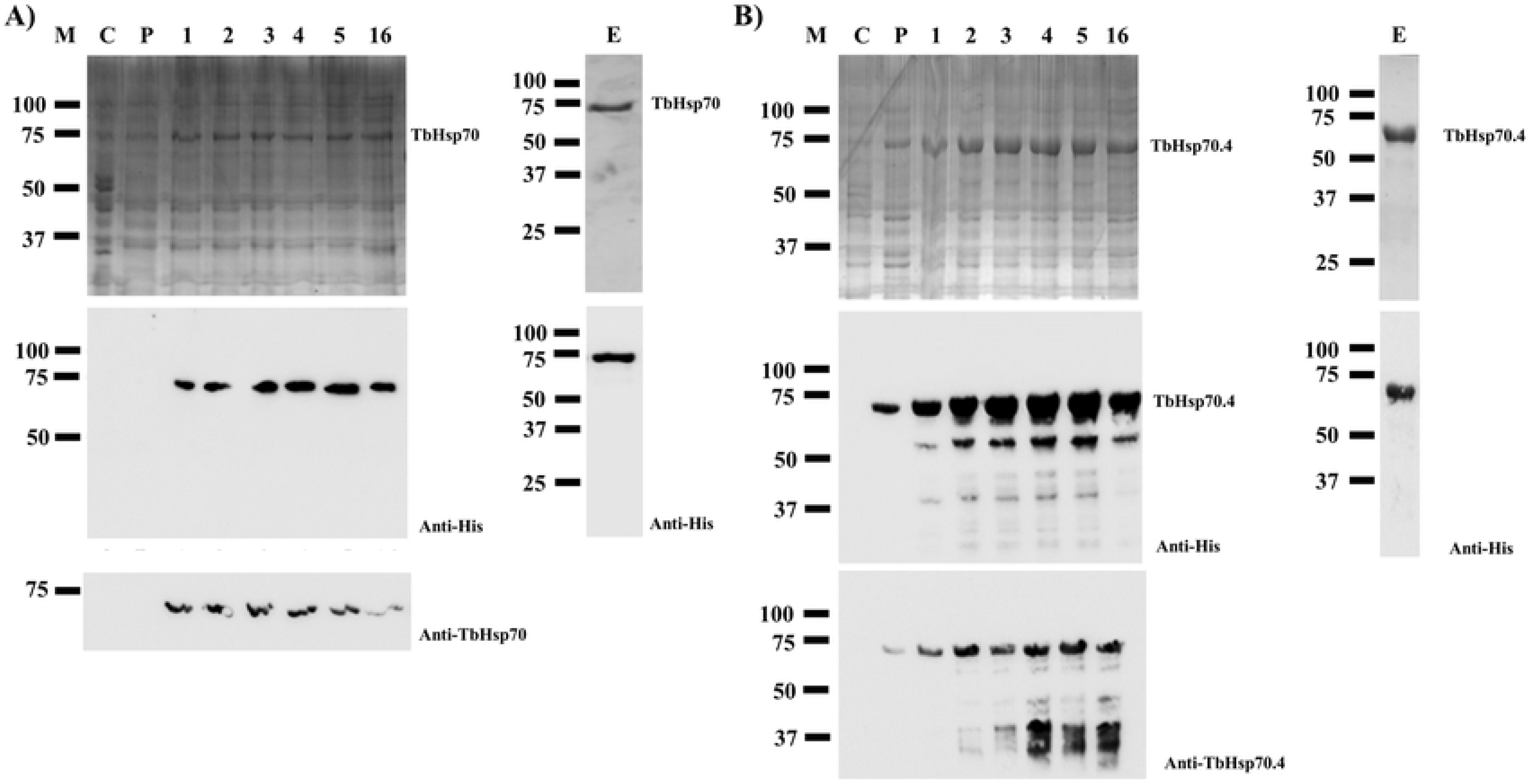
Expression and purification of recombinant TbHsp70 and TbHsp70.4. TbHsp70 and TbHsp70.4 were both expressed in *E. coli* XL1 Blue cells. SDS-PAGE (10%) and western blot images representing the expression and purification of recombinant forms of TbHsp70 (A) and TbHsp70.4 (B). *Lane M*: Marker in kilodalton (kDa) (Precision Plus Protein™ All Blue Prestained Protein Standard) is shown on the *left-hand side*; *Lane C*: The total extract for cells transformed with a neat pQE2 plasmid. *Lane P*: The total cell extract of *E. coli* XL1 Blue cells transformed with pQE2-TbHsp70, and pQE2-TbHsp70.4 prior to 1 mM IPTG induction; *Lane 1-16*: Total cell lysate obtained 1-16 h post induction, respectively. *Lane E*: Protein eluted from the affinity matrix using 500 mM imidazole. *Lower panels*: Western analysis using anti-His antibody to confirm expression and purification of recombinant TbHsp70 and TbHsp70.4. Western analysis using anti-TbHsp70 and anti-TbHsp70.4 to confirm expression of recombinant TbHsp70 and TbHsp70.4 were also used respectively.

### Protein expression of TbHsp70 and TbHsp70.4 is modulated in response to heat stress

The complex life cycle of *T. brucei* shows cell forms with different morphology and functional characteristics that interact with an insect vector and a mammalian host, undergoing several environmental variations in the process [32]. Hsp70 proteins have been shown in eukaryotic and prokaryotic cells to be self-protective proteins that maintain cell homeostasis against a wide variety of stressors as an adaptive response [33]. The impact of heat stress on TbHsp70 and TbHsp70.4 expression was assessed by exposing *T. b. brucei* 927 V221 parasites growing at the bloodstream stage to heat stress at 42 °C for one hour. Both Hsp70 proteins were shown to be expressed under normal culture conditions of 37°C (Fig 2A). The protein expression level of TbHsp70 was up-regulated in response to heat shock, whereas TbHsp70.4 was shown to be slightly down-regulated (Fig 2A-B). The differences in the protein levels observed for TbHsp70 and TbHsp70.4 for the various treatments were not due to differences in loading, as the levels of actin, the loading control, remained unchanged (Fig 2A, lower panel).

**Fig 2.**
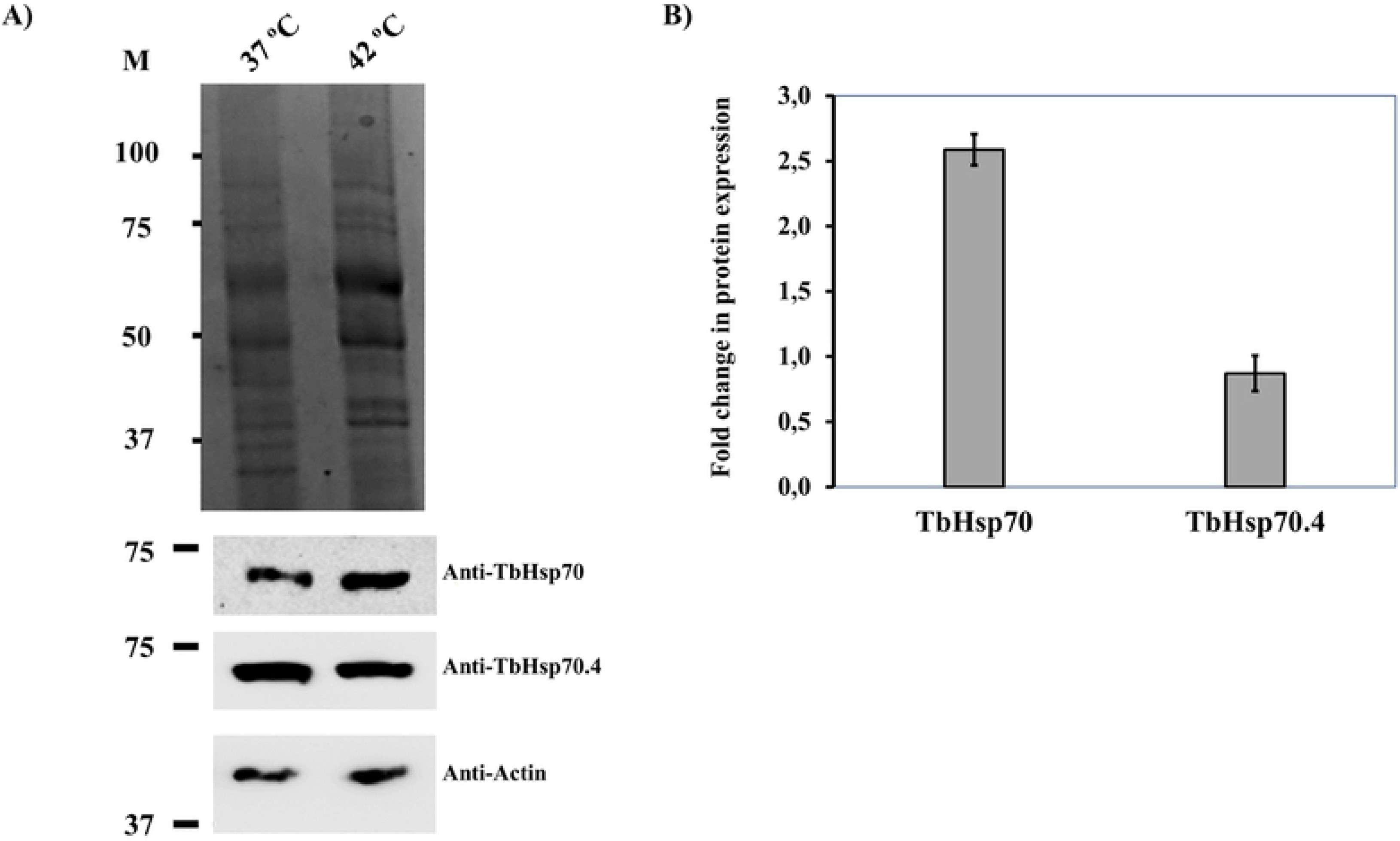
Expression of TbHsp70 and TbHsp70.4 in *T. b. brucei* parasites is modulated in response to heat stress at the blood stage. SDS-PAGE (10%) and western analyses of the expression of TbHsp70 and TbHsp70.4 by *T. b. brucei* parasites cultured at 37°C and 42°C, respectively (panel A); *Lane M* (molecular weight markers in kDa); parasite lysate harvested from bloodstream *T. b. brucei 927 V221* cells *in vitro* at 37°C and 42°C (5 × 106 cells/lane), respectively. *Lower panels:* Detection of the protein expression levels of TbHsp70 and TbHsp70.4 by western analysis using anti-TbHsp70 and anti-TbHsp70.4, respectively. Actin was used as a loading control to confirm that loading was equivalent in each lane. (B) Densitometric analysis illustrating the fold change in protein expression levels of the *T. brucei* Hsp70s in response to heat shock. A student’s t-test was used to validate the expression of the protein at 42°C compared to 37°C (p<0.005). The experiment was performed in triplicate using three different whole cell lysates and the figure represents the findings of a typical experiment.

The mRNA levels of TbHsp70 have been shown previously to be up-regulated in both transgenic bloodstream and procyclic form parasites in response to heat shock [34], as well as conditions that mimic mammalian infection [35]. TcHsp70, an orthologue of TbHsp70 in *Trypanosoma cruzi*, expression has also been investigated, and it has been shown that TcHsp70 synthesis increases 4-to 5-fold after heat shock [36], Thus, TbHsp70 is indicated to play a potential cytoprotective role in the parasite in response to various environmental stresses. The protein expression of TbHsp70.4 in response to heat stress is consistent with the findings observed for its orthologue in *Leishmania major* (LmjHsp70.4) [25]. The protein expression of LmHsp70.4 in promastigote parasites was shown to have no difference in expression after a period of heat stress [25]. In the cytoplasm of mammalian cells, the housekeeping functions are exerted by the constitutively expressed, cognate Hsp70 isoform, HSPA8 [37–38]. These functions include protein biogenesis, degradation, and protein translocation [8], which could infer that TbHsp70.4 may play a similar role in *T. brucei*. The modulation in protein expression of these Hsp70 proteins could indicate that Hsp70.4 and Hsp70 proteins in *T. brucei* and other kinetoplastid parasites may represent the cognate and inducible Hsp70 isoforms respectively.

### Localisation of TbHsp70 and TbHsp70.4

To further understand the biological role of the two Hsp70s, the sub-cellular localisation for TbHsp70 and TbHsp70.4 were investigated. The images for the sub-cellular localisation of the *T. brucei* Hsp70s, TbHsp70 and TbHsp70.4 were acquired from the TrypTag microscopy database [28]. This is a project that is aiming to tag every trypanosome protein with mNeonGreen (mNG) [39] in order to determine their localization in the cell [28]. Both TbHsp70 and TbHsp70.4 were C-terminally tagged with the mNG fluorescent protein. As depicted in the micrographs (Fig 3), both mNG-TbHsp70 and TbHsp70.4 were shown to localize in the cytosol of the parasite. The localization of TbHsp70.4 is consistent with the findings reported for its orthologue in *Leishmania major*, as the protein was shown through indirect immunofluorescence staining to reside in the cytosol [25]. The Hsp70 proteins also appeared to be excluded or depleted from the nucleus due to weaker fluorescence in those Hoechst-stained region (Fig 3). However, it has been demonstrated that the stress inducible Hsp70 proteins translocate from the cytosol to the nucleus in response to heat shock [40]. TcHsp70 has been shown to change its sub-cellular localisation from cytoplasmic to largely nuclear upon heat shock [41–42]. The Hsp70 isoform in the nucleus of *Trypanosoma cruzi* was shown to be different from its cytoplasmic counterpart indicating that the nuclear precursor protein must undergo specific modification prior to nuclear translocation [41]. TbHsp70 shares 89% sequence identity with TcHsp70, and therefore indicative that the protein may also be modified and change its sub-cellular localisation in response to heat shock or other environmental stresses. Thus, it would be important to further investigate the localization of TbHsp70 under stress conditions such as heat shock. TbHsp70 was also detected with high confidence from proteomic data to reside in the flagellum [43]. Inspection of the micrographs for TbHsp70-mNG illustrate punctuated flagellar localisation (Fig 3). TbHsp70.4-mNG also appears to reside in the flagellum but only at the posterior tip. Further investigation into flagellar localization is required, however, these findings may indicate that both *T. brucei* Hsp70s may also play a role in parasite mobility. The sub-cellular localization of Tbj2, was shown to reside in the cytosol of the parasite (data not shown), which is consistent with a previously reported localization for the J-protein [26]. The displayed cytosolic localization for both *T. brucei* Hsp70s and Tbj2 may suggest possible co-localization, and subsequently potential functional partnerships.

**Fig 3.**
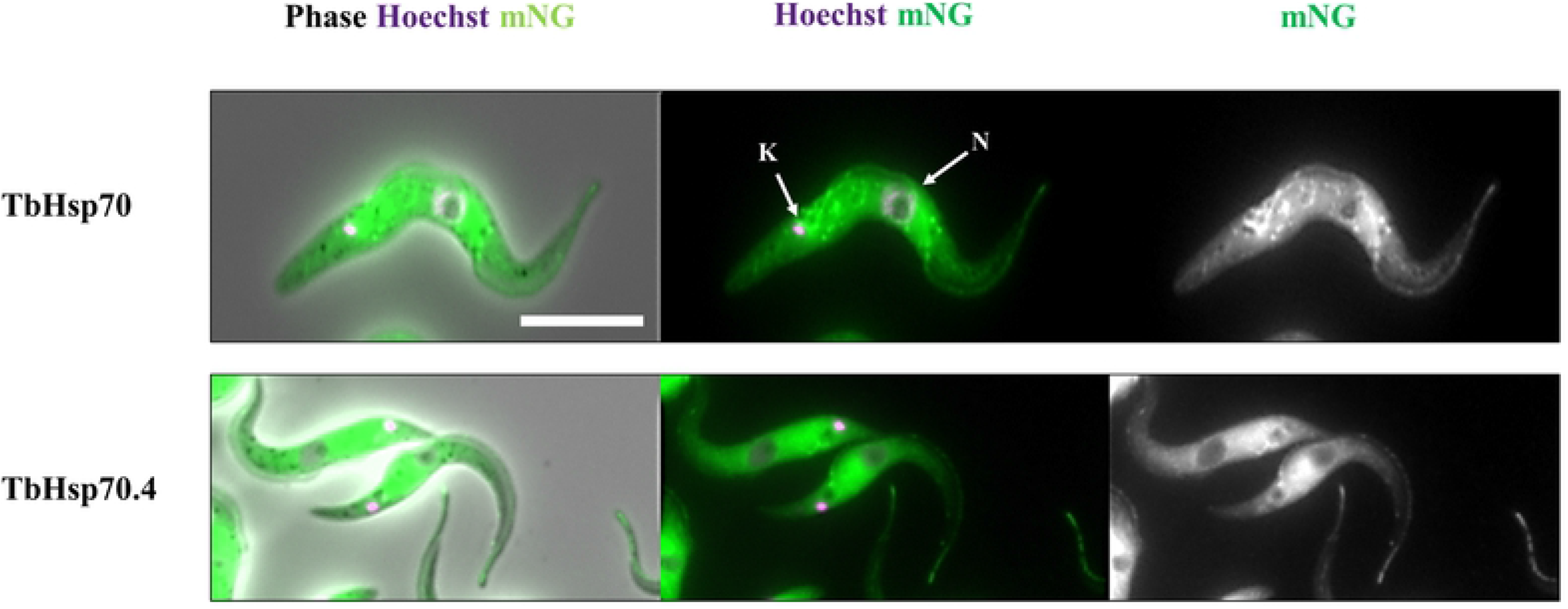
TbHsp70 and TbHsp70.4 are both localized to the parasite cytosol. Selected images from the TrypTag [28] high-throughput microscopy database was acquired for the investigation of the sub-cellular localisation of TbHsp70 (Tb927. 11.11330) and TbHsp70.4 (Tb927.7.710) in the parasite. Each protein was C-terminally tagged at the endogenous locus with the mNeonGreen (mNG) [39] fluorescent protein (*indicated as green*). Hoechst 33342, a fluorescent marker for DNA was used to stain both the nucleus (N) and kinetoplast (K) in the cell (*indicated as purple*). Different numbers of kinetoplasts and nuclei in the cells indicate different stages of the cell division cycle. The three panels from left to right display a representative image for the following: Phase contrast image of the merge and overlay of the mNG-tagged protein transfectants stained with Hoechst 33342; merge and overlay of the mNG-tagged protein transfectants stained with Hoechst 33342; distribution of mNG-tagged protein. Scale bar represents 5 μM.

### TbHsp70 and TbHsp70.4 both possess intrinsic ATPase activity which is stimulated by Tbj2

The basal ATPase activities for both TbHsp70 and TbHsp70.4 were determined using a colorimetric assay and were conducted under variable concentrations (0-2000 μM) of ATP. The Michaelis-Menten plots were generated from three independent batches of TbHsp70 and TbHsp70.4 (Fig 4). Both TbHsp70 and TbHsp70.4 exhibited intrinsic ATPase activity (Fig 4), though TbHsp70 was found to have a higher basal ATPase activity than TbHsp70.4 (Table S1) and the reported basal ATPase activity for TbHsp70.c (Table S1) [27]. However, the basal ATPase activities for the *T. brucei* Hsp70s were all found to be higher than those reported for human Hsp70 [42;44], bovine Hsc70 [45], and *E. coli* DnaK [46], but significantly lower than the values reported for Hsp70 homologues found in other protozoan parasites (Table S1). The K_m_ values obtained for TbHsp70 and TbHsp70.4, however, were found to be comparable indicating a similar affinity for ATP (Table S1).

**Fig 4.**
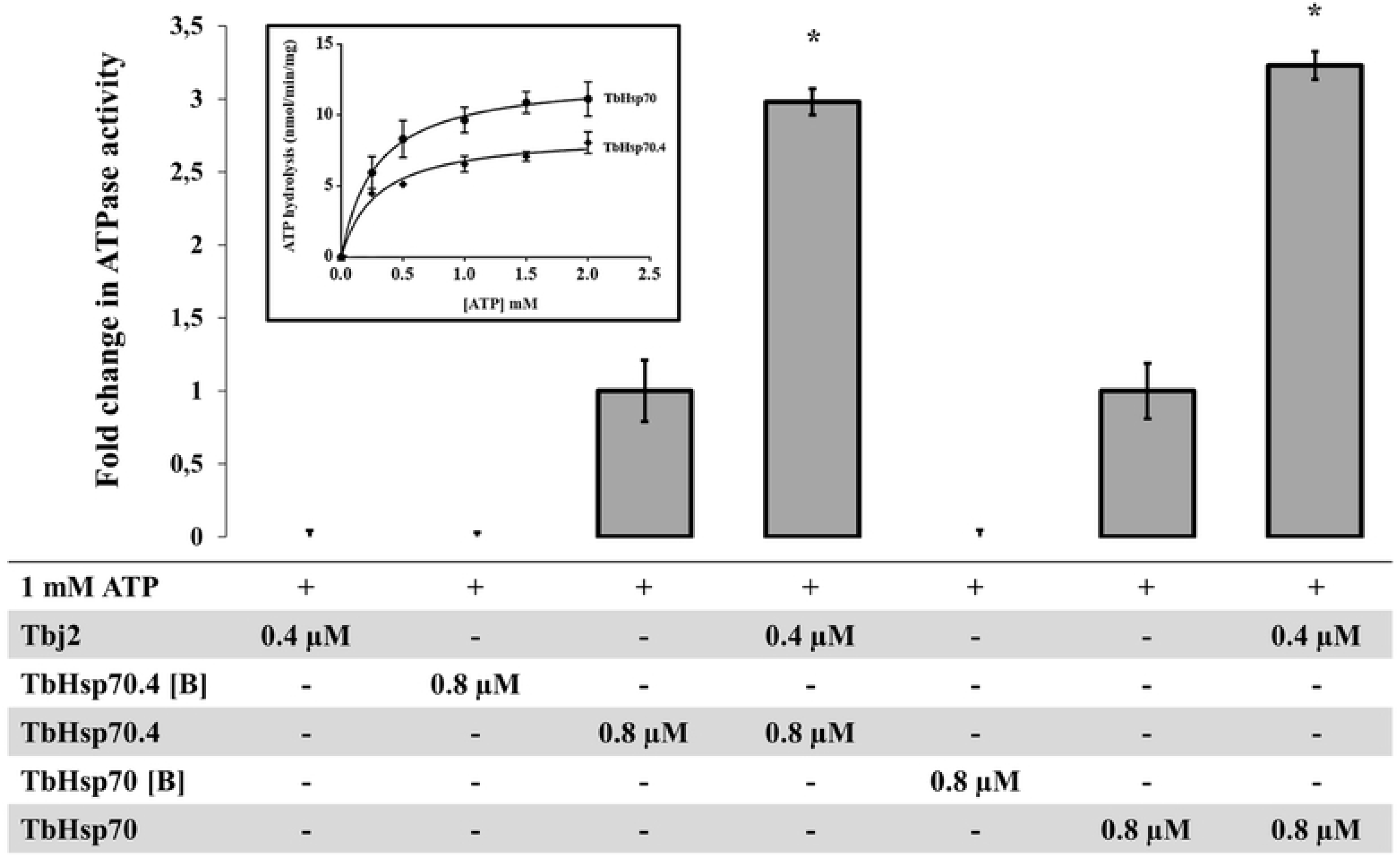
Stimulation of TbHsp70 and TbHsp70.4 ATPase activity by co-chaperone Tbj2. Submolar concentrations of recombinant Tbj2 were used to assess the ability of the co-chaperone to stimulate the basal ATPase activities of TbHsp70 and TbHsp70.4. The inorganic phosphate release was monitored by direct colorimetry at 595 nm wavelength. The curves represent the basal ATPase activities of recombinant TbHsp70 and TbHsp70.4, respectively expressed as mean (±) SD. The assay was carried out in triplicate from three independent batches of TbHsp70 and TbHsp70.4. ATP hydrolysis by the Tbj2 was also analysed separately and boiled samples of TbHsp70 and TbHsp70.4, indicated as [B] were included as negative controls. The ‘+’symbols represent components present in a reaction whilst ‘-‘represents those that were absent. The averaged data from three independent experiments done in triplicate using three independent batches of purified proteins for each experiment are shown with error bars indicated on each bar to represent standard deviation. Statistically significant difference of the ATPase activity of the TbHsp70s alone relative to the ATPase activity of the TbHsp70s in the presence of Tbj2 are indicated by * (P < 0.05) above the reaction.

As mentioned, the basal ATPase activity of Hsp70 is modulated by a cohort of co-chaperones, with the J-protein family being the most prominent. J-proteins stimulate the rate-limiting ATP hydrolysis step in the Hsp70 catalytic cycle, which facilitates substrate capture. Using a colorimetric assay, the effects of the cytosolic Type I J-protein from *T. brucei*, Tbj2, on the basal ATPase activity of TbHsp70 and TbHsp70.4 were explored. The basal ATPase activity of TbHsp70 and TbHsp70.4 was represented as 1, with modulation represented as a fold change in basal ATPase activity (Fig 4). Tbj2 was shown to moderately stimulate the basal ATPase activities of TbHsp70 and TbHsp704 by approximately 3 orders of magnitude (Fig 4). Neither Tbj2 nor TbHsp70 and TbHsp70.4 denatured by boiling displayed any ATPase activity (Fig 4) indicating that the increased phosphate release was due to stimulation of the Hsp70 ATPase activity. The magnitude of Hsp70 ATPase activity stimulation by Tbj2 is comparable to the study by Burger et al. [27] describing stimulation of TbHsp70.c and TcHsp70B by Tbj2. Thus, the results of these findings may indicate that Tbj2 may form a functional partnership with either of the cytosolic *T. brucei* Hsp70s.

### TbHsp70 and TbHsp70.4 both suppress the thermal aggregation of malate dehydrogenase (MDH)

The holdase function of TbHsp70 and TbHsp70.4 in the presence and absence of Tbj2 was determined by assessing its ability to prevent the thermally induced aggregation of the model substrate, malate dehydrogenase (MDH). The MDH aggregation suppression assay was adapted from [29]. MDH was subjected to heat stress at 48 °C and, as expected, the protein aggregated in the absence of chaperones (Fig 5). In the presence of BSA, a non-chaperone protein, MDH also aggregated in response to heat stress (Fig S2). In the absence of MDH, both *T. brucei* Hsp70s were found to be stable under the assay conditions as no significant aggregation was observed (Fig S2). The addition of either TbHsp70 or TbHsp70.4 at varying concentrations (0.25-0.75 μM) were both shown to suppress the thermally induced aggregation of MDH in a dose dependent manner (Fig S2). Though, TbHsp70 was shown to be more capable at suppressing the aggregation of MDH than TbHsp70.4 (Fig S2). TbHsp70 was shown to be heat inducible, and thus a more proficient holdase function maybe necessary for effective maintenance of the proteostasis under stressful conditions.

**Fig 5.**
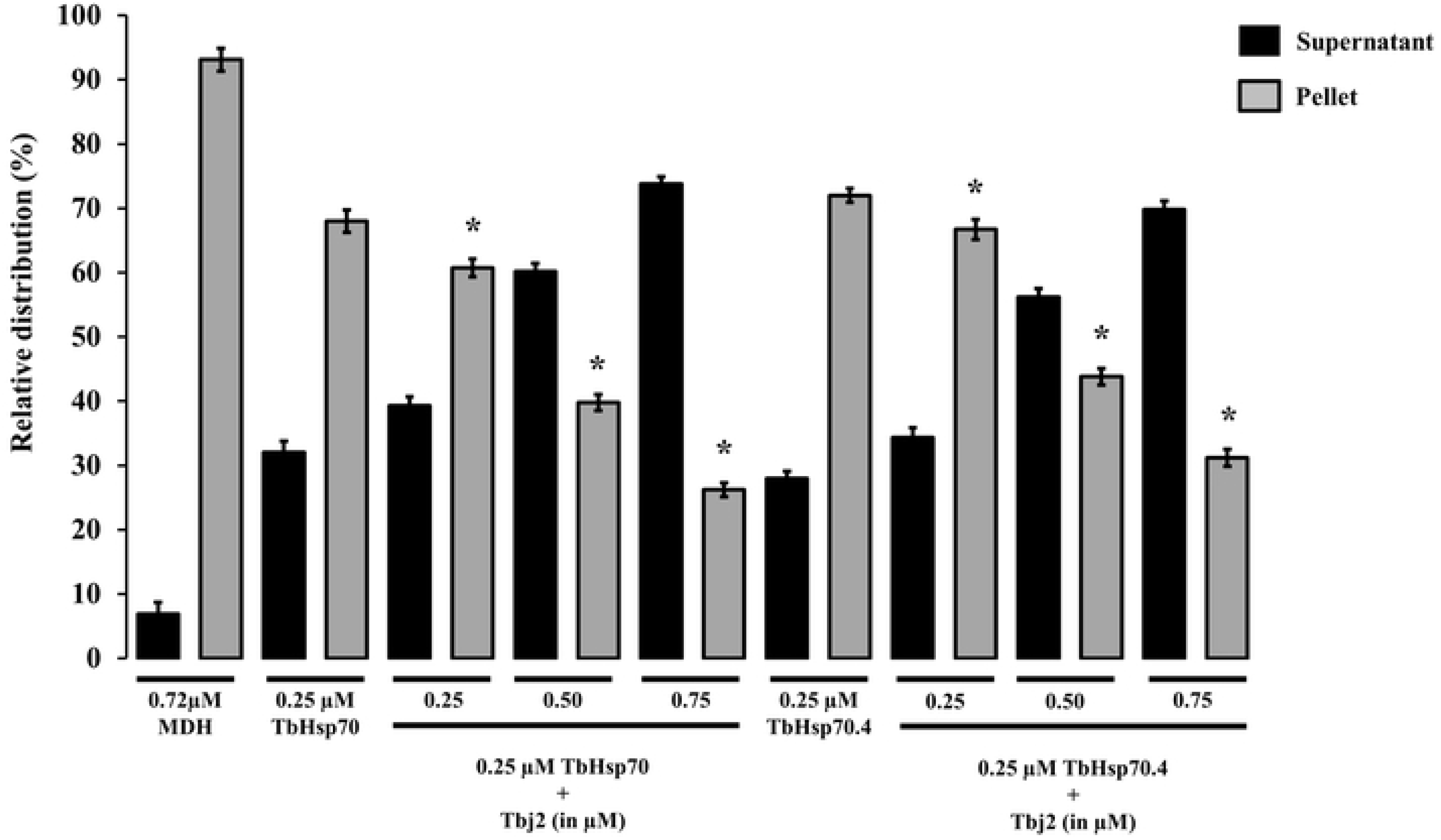
TbHsp70 and TbHsp70.4 suppresses the thermal aggregation of MDH. The holdase function of recombinant TbHsp70 and TbHsp70.4 was conducted by monitoring the heat-induced aggregation of MDH (0.72 μM) *in vitro* at 48°C and quantifying the pellet (insoluble; grey bars) and supernatant (soluble; black bars) fractions after heat exposure. The thermal aggregation of MDH in the presence of a non-chaperone, BSA is shown. TbHsp70.4 and TbHsp70 in the absence of MDH were not prone to aggregation under the assay conditions. Standard deviations were obtained from three replicate assays on three independent batches of purified protein. A statistically significant difference between a reaction and MDH alone are indicated by * (P < 0.05) above the reaction using a Student’s t-test.

The addition of Tbj2 at varying concentrations was also found to suppress the thermal aggregation of MDH (Fig S3), which is consistent with previous findings [27]. The Type I J-protein subfamily has been shown to possess intrinsic chaperone activity to select and deliver nascent polypeptides to their Hsp70 chaperone partner [47]. Tbj2 has been shown to be stress inducible [26] and the demonstrated ability to bind and suppress protein aggregates indicates that the J-protein may also play a prominent role in parasite cytoprotection. The ability of Tbj2 to enhance the holdase function of the *T. brucei* Hsp70s was assessed by maintaining constant Hsp70 concentration and varying those of the J-protein (Fig 5). TbHsp70 and TbHsp70.4 (0.25 μM) suppressed MDH aggregation suppression by 32% and 28% respectively (Fig 5). The addition of Tbj2 at equimolar concentration (0.25 μM) to TbHsp70 and TbHsp70.4 resulted in 39.3% and 34,3% MDH aggregation suppression respectively (Fig 5). Increasing concentrations of Tbj2 resulted in 60.2% (0.5 μM) and 73.8% (0.75 μM) MDH aggregation suppression for TbHsp70, and 56,2% (0.5 μM) and 69,8% (0.75 μM) MDH aggregation suppression for TbHsp70.4 (Fig 5). Despite Tbj2 being shown to possess independent chaperone activity (Fig S3), the modulatory effect of Tbj2 on the holdase activity of the *T. brucei* Hsp70s was not found to be additive, indicating a demonstrated synergistic effect and thus prompting Tbj2 as a potential co-chaperone to both TbHsp70 and TbHsp70.4.

### Tbj2 stimulates the refoldase activities of TbHsp70 and TbHsp70.4

A β-galactosidase refolding assay, adapted from [30], was used to investigate the refoldase activity of TbHsp70 and TbHsp70.4 in the presence and absence of Tbj2. β-galactosidase recognises and cleaves the chromogenic substrate, o-nitrophenyl-β-D-galactoside (ONPG), into ortho-nitrophenol (ONP) and galactose. Absorbance at 420 nm measures the amount of ONP produced in the reaction which reflects the ability of the molecular chaperones to refold chemically denatured β-galactosidase. In these experiments, the activity of β-galactosidase without denaturation was considered as 100%. It was shown that the *T. brucei* Hsp70s were not able to proficiently mediate β-galactosidase refolding (Fig 6). Human Hsp70 has also been demonstrated unable to independently mediate the refolding of chemically denatured β-galactosidase [30] or thermally denatured luciferase [48]. It is suggested that the Hsp70 preferentially maintains the non-native substrate in a ‘folding-competent’ state which, upon addition of a J-protein co-chaperone, leads to refolding of the client protein [30]. The addition of Tbj2 to TbHsp70 or TbHsp70.4 resulted in the stimulation of the refoldase activity of the Hsp70s, and approximately 50% of the non-native β-galactosidase being refolded (Fig 6). These findings, however, are comparable with those reported for the refolding of non-native β-galactosidase mediated by human Hsp70 and the Type I J-protein, Hdj1, which was shown to reactivate approximately 40% of the denatured enzyme [30]. However, the stimulation of the refoldase activity of both TbHsp70 and TbHsp70.4 by Tbj2 indicates the formation of a functional Hsp70/J-protein partnership.

**Fig 6.**
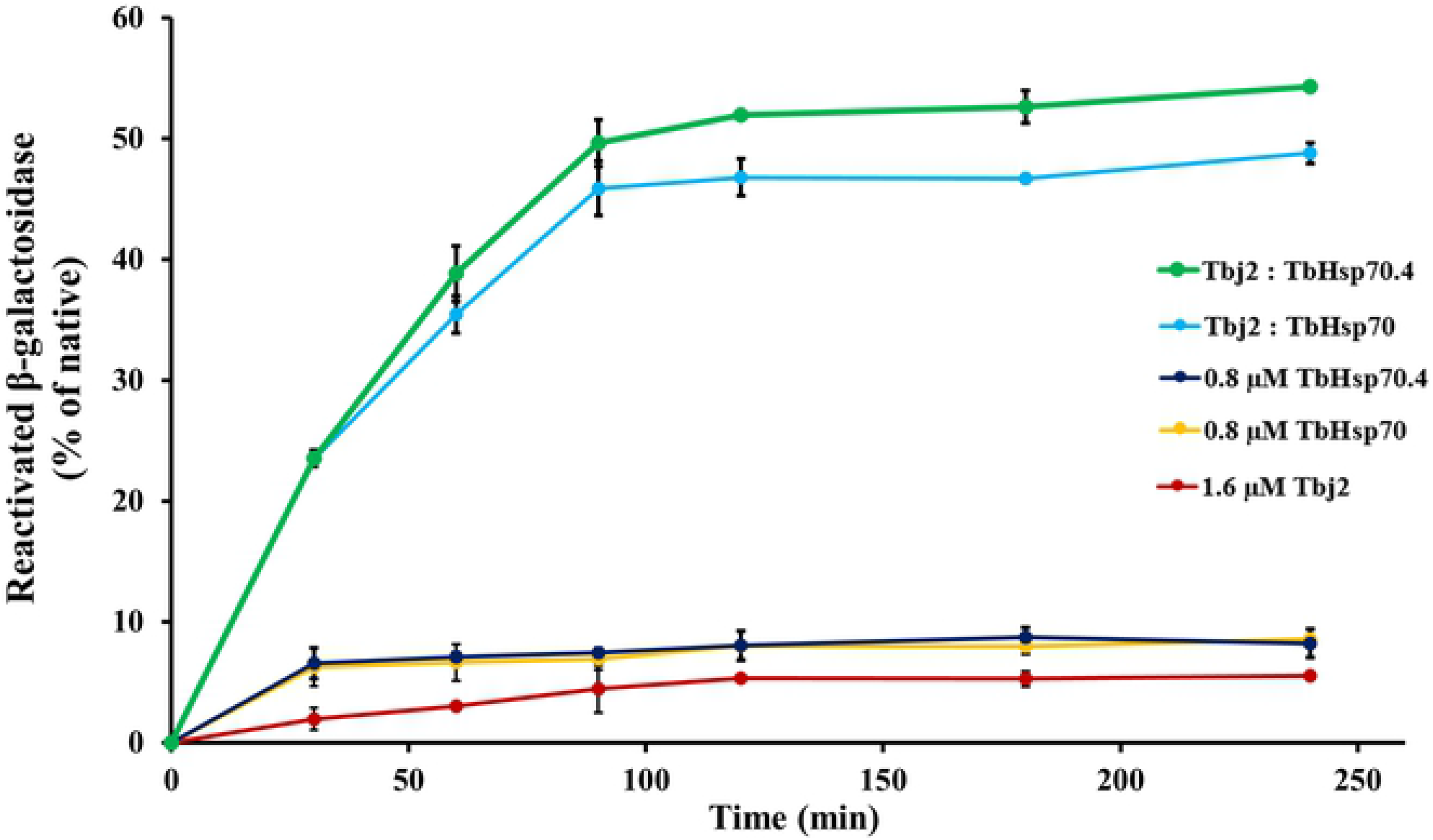
Tbj2 stimulates the refoldase activity of TbHsp70 and TbHsp70.4. Chemically-denatured β-galactosidase (3.4 nM) was diluted into refolding buffer containing the indicated Hsp70 and/or Tbj2 mixtures. β-galactosidase assays were performed as described [30]. β-galactosidase activity was measured using ONPG as a chromogenic substrate. Results are expressed as % β-galactosidase activity of the refolded enzyme in relation to native enzyme. Standard deviations were obtained from three replicate assays on three independent batches of purified protein.

## Conclusion

It has become increasingly evident that the Hsp70/J-protein machinery is essential to the survival, pathogenicity, and differentiation of kinetoplastid parasites. However, further elucidation of the molecular details of the Hsp70/J-protein chaperone interactions and pathways are required, as some of these pathways may represent a novel means of chemotherapeutic intervention for African trypanosomiasis. This study aimed to investigate the cytosolic Hsp70 system, through *in vitro* biochemical characterization of the chaperone properties of the predicted cytosolic *T. brucei* Hsp70s, TbHsp70 and TbHsp70.4, and their potential partnership with the cytosolic Type I J-protein, Tbj2. We have provided the first evidence that TbHsp70 and TbHsp70.4 represent the heat-inducible and cognate Hsp70 isoforms, respectively, that reside in the cytosol of the parasite. Thus, TbHsp70 may represent an essential component in cytoprotection of parasite, whereas TbHsp70.4 fulfils crucial housekeeping roles. Tbj2 was shown able to stimulate the basal ATPase activity of both Hsp70s and form a functional partnership capable of mediating the reactivation of client proteins. Overall, this study provides a greater understanding of the *T. brucei* Hsp70 chaperone system, and the critical role these proteins play in the cell biology of the parasite. Further studies, however, are required to explore the *T. brucei* Hsp70/J-protein machinery, and to identify small-molecule inhibitors capable of disrupting these systems.

## Acknowledgements

The pET28a-Tbj2 and pQE2-TbHsp70.4 expression vectors were kindly provided by Dr Michael Ludewig (Rhodes University, South Africa). The *T. b. brucei* Lister 927 variant 221 strain was a kind donation from Professor George Cross (Rockefeller University, New York, U.S.A.). Ms Michelle Isaacs (Rhodes University) for *T. b. brucei* culturing at the parasite facility, which is supported by the South African Medical Research Council (MRC). The rabbit polyclonal anti-TbHsp70 and rabbit polyclonal anti-TbHsp70.4 antibodies were generous gifts from Dr Ariel Louwrier (StressMarq Biosciences Inc., Canada) and Professor Jay Bangs (University of Buffalo, New York, U.S.A.) respectively. We would also like to acknowledge Richard Wheeler from the TrypTag high-throughput microscopy database (TrypTag.org) for the localisation images.

## Funding

This work was funded by a grant from the National Research Foundation (NRF), grant number 87663. S.J.B. was a recipient of an NRF Doctoral Innovation Scholarship.

## Supporting Information

**Table S1. Kinetic constants determined for TbHsp70 and TbHsp70.4 compared with those of homologous Hsp70.**

**Fig S1. Expression and purification of recombinant Tbj2.** Recombinant Tbj2 was expressed in *E. coli* BL21 (DE3) cells. SDS-PAGE (10%) and western blot images representing the expression (A) and purification (B) of recombinant forms of Tbj2. *Lane M*: Marker in kilodalton (kDa) (Precision Plus Protein™ All Blue Prestained Protein Standard) is shown on the left-hand side; *Lane C*: The total extract for cells transformed with a neat pET28a plasmid. *Lane P*: The total cell extract of *E. coli* BL21 (DE3) cells transformed with pET28a-Tbj2 prior to 1 mM IPTG induction; *Lane 1-16*: Total cell lysate obtained 1-16 h post induction, respectively. *Lane E*: Protein eluted from the Ni2+ chelate affinity matrix using 500 mM imidazole. *Lower panels*: Western blots on use of anti-His antibody to confirm expression and purification of recombinant Tbj2.

**Fig S2. TbHsp70 and TbHsp70.4 suppresses the thermal aggregation of MDH.** The holdase function of recombinant TbHsp70 and TbHsp70.4 was conducted by monitoring the heat-induced aggregation of MDH (0.72 μM) in vitro at 48°C and quantifying the pellet (insoluble; grey bars) and supernatant (soluble; black bars) fractions after heat exposure. The thermal aggregation of MDH in the presence of a non-chaperone, BSA is shown. TbHsp70.4 and TbHsp70 in the absence of MDH were not prone to aggregation under the assay conditions. Standard deviations were obtained from three replicate assays on three independent batches of purified protein. A statistically significant difference between a reaction and MDH alone are indicated by * (P < 0.05) above the reaction using a Student’s t-test.

**Fig S3: Tbj2 suppresses the thermal aggregation of MDH.** The chaperone activity of Tbj2 was conducted by monitoring the heat-induced aggregation of MDH (0.72 μM) *in vitro* at 48°C and quantifying the pellet (insoluble; grey bars) and supernatant (soluble; black bars) fractions after heat exposure. Tbj2 in the absence of MDH was not prone to aggregation under the assay conditions. Standard deviations were obtained from three replicate assays on three independent batches of protein. A statistically significant difference between a reaction and MDH alone are indicated by * (P < 0.05) above the reaction using a Student’s t-test.

